# The structural basis for Ulp2 recruitment to the kinetochore

**DOI:** 10.1101/2020.12.30.424835

**Authors:** Yun Quan, Stephen M. Hinshaw, Pang-Che Wang, Stephen C. Harrison, Huilin Zhou

## Abstract

The step-by-step process of chromosome segregation defines the stages of the cell division cycle. In eukaryotes, signaling pathways that control these steps converge upon the kinetochore, a multiprotein assembly that connects spindle microtubules to the centromere of each chromosome. Kinetochores control and adapt to major chromosomal transactions, including replication of centromeric DNA, biorientation of sister centromeres on the metaphase spindle, and transit of sister chromatids into daughter cells during anaphase. Although the mechanisms that ensure tight microtubule coupling at anaphase are at least partly understood, kinetochore adaptations that support other cell cycle transitions are not. We report here a mechanism that enables regulated control of kinetochore sumoylation. A conserved surface of the Ctf3/CENP-I kinetochore protein provides a binding site for the SUMO protease, Ulp2. Ctf3 mutations that disable Ulp2 recruitment cause elevated inner kinetochore sumoylation and defective chromosome segregation. The location of the site within the assembled kinetochore suggests coordination between sumoylation and other cell cycle-regulated processes.

## INTRODUCTION

The cellular challenge of accomplishing error-free chromosome segregation is a central problem in biology. Success is essential for the health and longevity of all multicellular organisms. Environmental variability ensures that no two cell divisions are identical, and cells modulate the activities of chromosome segregation machines accordingly. One classic example of such regulation is the mitotic checkpoint, which encompasses a collection of kinases and associated scaffold proteins that prevent progression to anaphase until all pairs of sister centromeres are properly aligned on the mitotic spindle (London and Biggins, 2014). In addition to phosphorylation cascades, cells use the <u>S</u>mall <u>U</u>biquitin-like <u>M</u>odifier (SUMO) protein to ensure high-fidelity chromosome segregation (Fukagawa et al., 2001; Li and Hochstrasser, 2000; Meluh and Koshland, 1995). One important mechanism involves regulated sumoylation of distinct sets of kinetochore substrates (de Albuquerque et al., 2016; Montpetit et al., 2006; Mukhopadhyay et al., 2010). Mechanisms that target SUMO pathway components to these kinetochore subsets have not been fully described. Doing so is a prerequisite for understanding their regulation in response to environmental and cell cycle cues.

Yeast kinetochores, which are thought to be simplified versions of larger vertebrate kinetochores, contain two functional domains (Biggins, 2013) that can be further subdivided into biochemically defined multiprotein complexes (Cheeseman et al., 2002; De Wulf et al., 2003; Hinshaw and Harrison, 2018; McKinley and Cheeseman, 2016; Musacchio and Desai, 2017). Outer kinetochore proteins contact microtubules and are the principal substrates and organizers of the mitotic checkpoint. Inner kinetochore proteins, most of which assemble into the Ctf19 complex (Ctf19c in yeast, <u>C</u>onstitutive <u>C</u>entromere <u>A</u>ssociated <u>N</u>etwork or CCAN in vertebrates), contact centromeric DNA and regulate chromosomal functions. Structural studies of inner kinetochore proteins imply regulated chromatin recognition and show how key activities are coordinated (Hinshaw and Harrison, 2019, 2020; Kixmoeller et al., 2020; Yan et al., 2019).

SUMO pathway components were identified due to their ability, when overexpressed, to rescue the viability of lethal kinetochore mutants (Meluh and Koshland, 1995). One such factor, the nuclear SUMO protease Ulp2 (homologous to human SENP6), cleaves SUMO chains from substrate proteins (Li and Hochstrasser, 2000). Ulp2 activity coordinates multiple chromosomal functions, and its localization and substrate recognition are the main points of regulation (Kroetz and Hochstrasser, 2009). Differential targeting depends on short peptide motifs embedded within N- and C-terminal extensions flanking a central catalytic domain. In particular, Ulp2 residues 781-873 contact Csm1, a component of the monopolin complex (de Albuquerque et al., 2018; Liang et al., 2017), and Ulp2 residues 896-937 (previously CCR for <u>C</u>onserved <u>C</u>-terminal <u>R</u>egion, and renamed here KIM for <u>K</u>inetochore <u>I</u>nteraction <u>M</u>otif) contact the inner kinetochore Ctf3 complex (Ctf3c, CENP-H/I/K in vertebrates) (Suhandynata et al., 2019). A <u>S</u>UMO-<u>I</u>nteracting <u>M</u>otif (SIM) comprising Ulp2 residues 725-728 boosts Ctf3c- and Csm1-dependent Ulp2 activity at kinetochores and the nucleolus, respectively (de Albuquerque et al., 2018; Suhandynata et al., 2019). SIM-dependence implies homeostatic regulation; excessive substrate sumoylation enhances Ulp2 recruitment, and cleavage of the chains by Ulp2 itself releases the enzyme from its substrates.

Sumoylated inner kinetochore proteins accumulate in cells expressing Ulp2 kinetochore interaction mutants (Ulp2-SIM-3A-KIM-3A) (Suhandynata et al., 2019). These cells, like *ulp2*Δ cells, frequently missegregate chromosomes (Ryu et al., 2016; Suhandynata et al., 2019). Likewise, human SENP6 cleaves SUMO from inner kinetochore proteins (Fu et al., 2019; Liebelt et al., 2019; Mitra et al., 2020; Mukhopadhyay et al., 2010; Wagner et al., 2019), and SENP6 depletion destabilizes kinetochores, causing severe chromosome segregation defects (Liebelt et al., 2019; Mitra et al., 2020; Mukhopadhyay et al., 2010). These findings suggest a conserved mechanism that counteracts inner kinetochore sumoylation and maintains kinetochore integrity. Identification of a kinetochore receptor for Ulp2 supports this viewpoint (Suhandynata et al., 2019), but both the receptor (the Ctf3c) and Ulp2 itself have multiple functions. A definitive account of the contribution of kinetochore-directed Ulp2 activity to accurate chromosome segregation and related studies of cell cycle-regulated Ulp2 activity in this process thus require a finer description of Ulp2 recruitment.

We present here an analysis of the kinetochore targeting function of Ulp2. A structure of the Ctf3c bound to the Ulp2-KIM shows that Ulp2 contacts a conserved surface of the Ctf3 protein. Mutation of the Ctf3 surface causes elevated inner kinetochore sumoylation. The phenotype is especially pronounced in strains that also carry the *ulp2-SIM-3A* mutation, and the ultimate consequence of this dysregulation is defective chromosome segregation in the mutant strains. These findings provide a conclusive demonstration of targeted kinetochore desumoylation by Ulp2 and suggest mechanisms that might regulate this activity during the cell cycle.

## RESULTS

### The structure of the Ctf3c interacting with Ulp2

To determine how the Ctf3c recruits Ulp2 to the kinetochore, we determined the structure of the Ctf3c-Ulp2 complex by single particle cryo-EM. We first reconstituted the interaction between purified Ulp2-KIM and the Ctf3c (Figure 1B). The previously described Ulp2-KIM-3A mutant (Ulp2-V931A,L933A,I934A) did not bind the Ctf3c, nor did the SIM that bolsters the kinetochore recruitment of the enzyme (Ulp2 residues 725-728, Figure S1A), demonstrating the specificity of the interaction. A minimal Ulp2 peptide (Ulp2 residues 927-937) bound the Ctf3c in a pulldown assay with an affinity that matched longer Ulp2 fragments (Figure S1B). We purified a complex containing this minimal Ulp2 peptide and the Ctf3c (Figure 1C) and determined its structure by cryo-EM (Figure 1D and S1C).

**Figure 1.**
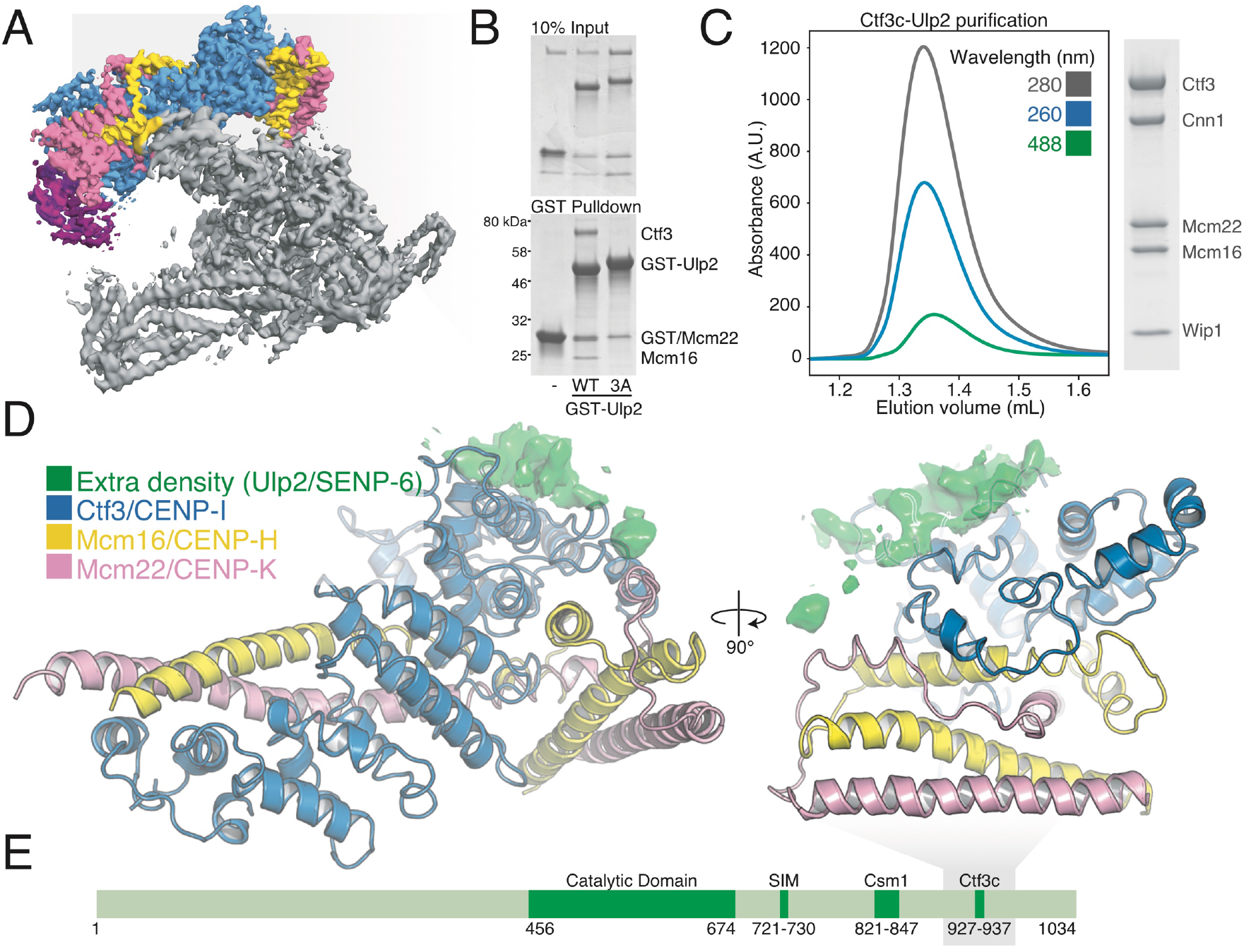
Reconstitution and structure of the Ctf3c-Ulp2 complex. A) Model of the assembled Ctf19c (Ctf3 – blue; Mcm22 – pink; Mcm16 – yellow; Cnn1/Wip – purple; Chl4, Iml3, Ctf19, Mcm21, Ame1, Okp1, Nkp1, Nkp2 – gray; EMD-0523, EMD-2910) (Hinshaw and Harrison, 2019, 2020). B) The Ctf3c binds the Ulp2-KIM but not the Ulp2-KIM-3A peptide. Purified Ctf3c and GST-Ulp2-KIM were mixed and subjected to pulldown on glutathione beads. Input and pulldown samples were analyzed by SDS-PAGE. C) Purification of the Ctf3c-Ulp2-KIM complex by size exclusion chromatography. Absorbance wavelengths are indicated. 488 nm reports on the FITC-Ulp2-KIM peptide. Eluate from the peak fractions were analyzed by SDS-PAGE. D) Structure of the Ctf3c-Ulp2 complex. Extra density corresponding to Ulp2-KIM is shown as a green surface derived from the final density map. E) Domain diagram of Ulp2. Regions with known functions are annotated.

The Ulp2-Ctf3c structure shows the Ulp2-KIM interacting with the C-terminal region of the Ctf3 protein. Density that was not part of the Ctf3c marked the position and contact surface of the Ulp2-KIM peptide but was not sufficiently well defined to permit assignment of individual residue positions. The full Ctf3c comprises two structural modules, defined by N- and C-terminal Ctf3 HEAT repeat arrays that contact parallel coiled-coils belonging to the Mcm16 and Mcm22 proteins (Hinshaw and Harrison, 2020). The Ulp2-KIM contacts the exterior surface of the Ctf3 C-terminal HEAT array near its C-terminus and does not interact with either Mcm16 or Mcm22. In the context of the complete Ctf3c, the Ulp2 interaction surface is exposed to solvent.

Previous cryo-EM structures of the intact Ctf19c enable analysis of the Ulp2 binding site relative to other inner kinetochore proteins (Hinshaw and Harrison, 2019; Yan et al., 2019). The Ulp2-binding surface of the Ctf3c faces solvent in both published Ctf19c structures, consistent with the finding that Ulp2 localizes to centromeres and acts on assembled kinetochores. Cnn1-Wip1, histone fold proteins that bind Ctf3-N and indirectly connect the inner kinetochore to spindle microtubules, are distal to the Ulp2-KIM site. The arrangement implies a functional specialization of the Ctf3-N and −C modules whereby Ctf3-N mediates microtubule contact through Cnn1, and Ctf3-C regulates chromatin-associated processes through Ulp2. The Ulp2-KIM binds the Ctf3c near the anchor points for extended N-terminal regions of the Ctf19 and Mcm21 proteins, members of the COMA complex. These extended peptides are likely sites of post-translational modifications by SUMO ligases, indicating a mechanism for multi-site recognition of kinetochore architecture by Ulp2 through both its KIM and SIM peptides (Figure 1E).

### Analysis of the Ulp2-Ctf3c interaction: specificity and affinity

The Ctf3 surface that binds Ulp2 is conserved, suggesting its function in kinetochore regulation may have been retained throughout evolution (Figure 2A). To verify the position of the Ulp2-KIM relative to Ctf3, we mutated two conserved basic amino acid residues in Ctf3 to create the Ctf3-2A protein (Ctf3-R594A,K596A). Pulldown assays showed that Ctf3-2A does not support the Ulp2-KIM interaction (Figure S2A). The dissociation constant for the Ulp2-KIM-Ctf3c interaction, measured by fluorescence polarization using a minimal Ulp2 peptide, is ~0.9 μM (Figure 2B). The affinity of the same peptide for a Ctf3c containing the Ctf3-2A protein was not measurable. Unlabeled recombinant Ulp2-KIM inhibited the interaction with wild type Ctf3c, whereas Ulp2-KIM-3A did not, demonstrating the specificity of the assay.

**Figure 2.**
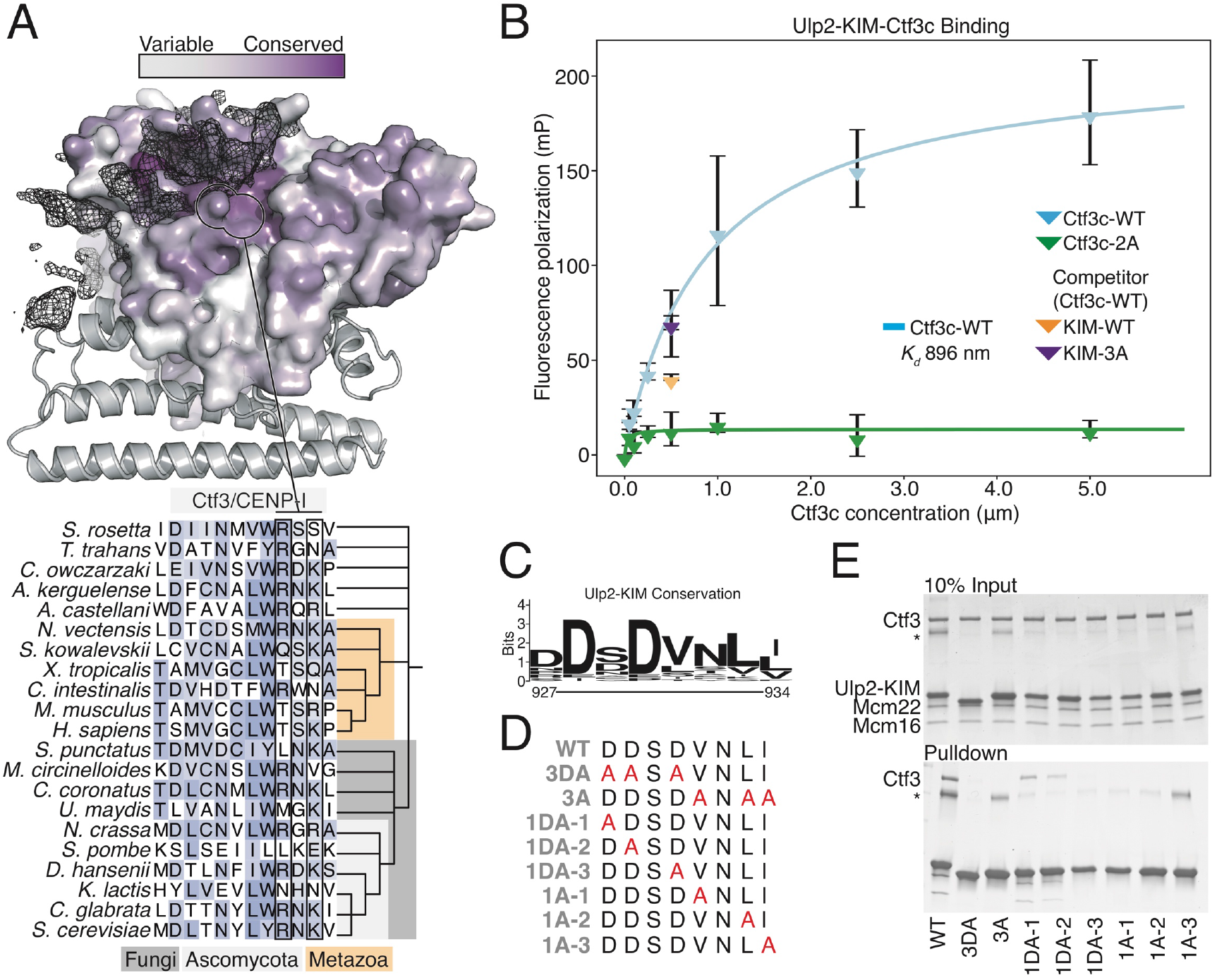
Biochemical analysis of the Ctf3c-Ulp2 interaction. A) Structure of the Ctf3c showing the Ctf3 protein surface colored according to Ctf3/CENP-I amino acid conservation. Mcm16 and Mcm22 are shown in cartoon view in gray. Extra density corresponding to the Ulp2-KIM is shown as a mesh. The alignment below shows Ctf3/CENP-I homologs from the indicated species. B) Ulp2-KIM binds the Ctf3c with a dissociation constant of ~0.9 μM. Fluorescence polarization (Y-axis) was measured for varying concentrations (X-axis) of purified Ctf3c (blue) or Ctf3c-2A (green). Three measurements for each Ctf3c concentration were averaged and plotted (error bars – +/- SD). C) Sequence logo showing the conservation of Ulp2-KIM residues across budding yeasts. D) Catalog of the Ulp2-KIM peptides used in panel E. E) Single amino acid substitutions in the Ulp2-KIM disrupt Ctf3c binding. Glutathione pulldown was performed to detect Ctf3c binding to GST-Ulp2-KIM with the indicated substitutions (* contaminant).

Clusters of basic and non-polar amino acid side chains dominate the Ctf3 conserved surface. Basic Ctf3 amino acid side chains likely satisfy the conserved negatively charged KIM aspartates (Ulp2-D928,D930), and non-polar Ctf3 residues likely provide a docking site for the conserved non-polar residues mutated in the Ulp2-KIM-3A protein (Figure 2D). To test this idea, we performed binding experiments with mutated minimal Ulp2-KIM proteins to probe the Ctf3c interaction (Figure 2E). Mutagenesis of conserved Ulp2 aspartate residues (Ulp2-3DA, 1DA-2, and 1DA-3) to alanine produced KIM peptides that did not interact with Ctf3. Mutation of Ulp2-D927 (1DA-1), which is less conserved, had only a modest effect. The Ulp2-KIM-3A peptide and related variants with single conserved hydrophobic KIM residues mutated to alanine (Ulp2-1A-1, 1A-2, and 1A-3) did not bind the Ctf3c. A Ulp2-KIM peptide bearing the N932A mutation (Figure S2B) retained Ctf3c binding activity. These findings coincide with conservation among fungal Ulp2-KIM peptides and support the mode of binding indicated by structural studies of the complex.

### The ctf3-2A mutation increases kinetochore sumoylation

To determine whether the Ctf3-Ulp2 interaction identified in our structural and biochemical studies dictates Ulp2 kinetochore recruitment *in vivo*, we introduced the *ctf3-2A* allele into the chromosomal *CTF3* locus. Isolation of Ulp2-KIM-binding proteins from cell extracts confirmed that Ctf3 but not Ctf3-2A binds Ulp2 (Figure 3A). The effect was even more pronounced when the Ulp2-KIM-3A mutant peptide was used as bait, consistent with a mode of interaction that depends on both polar and nonpolar contacts, as described above. To test whether the impaired Ulp2-Ctf3-2A interaction disrupts Ulp2 recruitment to centromeres, we analyzed Ulp2 localization by ChIP-qPCR (Figure 3B). Ulp2 is detectable above background at centromeres (*CEN3*) in wild type cells, and this localization is disrupted to similar degrees in *ulp2-KIM-3A, ctf3-2A*, and double-mutant (*ulp2-KIM-3A ctf3-2A*) cells. Therefore, the conserved surface of Ctf3 is required for Ulp2 recruitment to the centromere.

**Figure 3.**
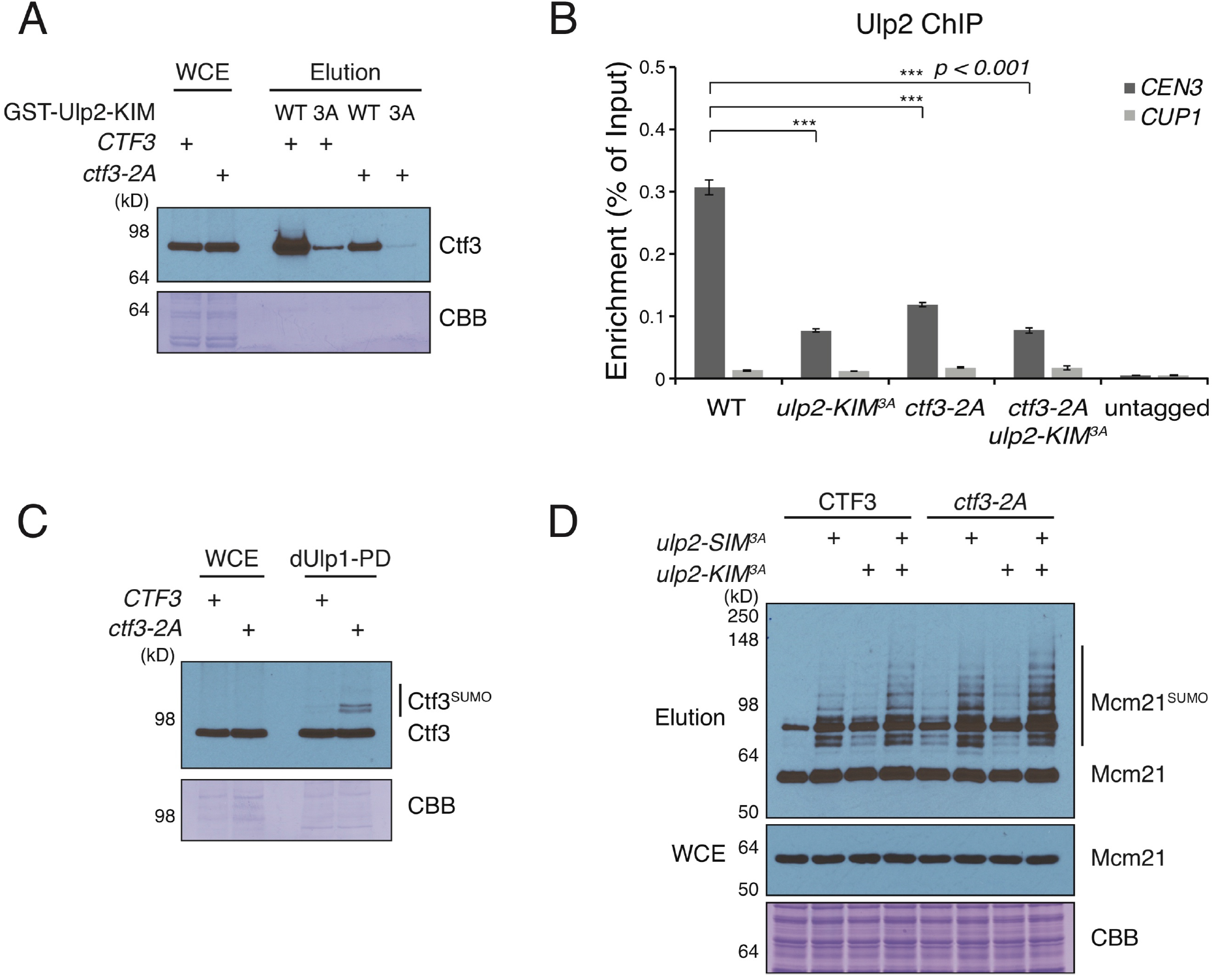
Ctf3 mutations that disrupt Ulp2 binding cause elevated kinetochore sumoylation. A) *ctf3-2A* reduces the amount of Ctf3 pulled down from yeast cell extracts by recombinant GST-Ulp2-KIM, and this interaction is further reduced by *ulp2-KIM-3A* mutation. Immobilized GST-Ulp2-KIM protein (wild type or KIM-3A as indicated; WCE – whole cell extract; CBB – Coomassie brilliant blue) was used as bait for affinity purification of Ctf3 from cells expressing either Ctf3 or Ctf3-2A. Tagged Ctf3 from cell lysate was detected using an anti-protein A antibody. B) Ulp2 centromere recruitment is defective in *ctf3-2A* and *ulp2-KIM-3A* cells. ChIP-qPCR was used to evaluate the effect of *ctf3-2A* on Ulp2 localization at a centromeric (*CEN3*) and a non-centromeric locus (*CUP1*) (*** – *p* ≤ 0.001; Student’s t-test, two tails, equal variance). C) and D) Effect of *ctf3-2A* and *ulp2-SIM-3A* on sumoylation of Ctf3 and Mcm21. Total sumoylated proteins from each indicated strain were purified using dUlp1 affinity resin and analyzed by antiprotein A Western blotting.

We next asked whether impaired Ulp2-Ctf3c binding results in elevated sumoylation of inner kinetochore proteins. To do so, we used immobilized, catalytically inactive Ulp1 (dUlp1) to isolate sumoylated cellular proteins and their complexes from cell extract. We observed the appearance of a prominent new species corresponding to sumoylated Ctf3 in the SUMO-enriched material from *ctf3-2A* cells (Figure 3C). Therefore, the inability of Ulp2 to bind kinetochores in *ctf3-2A* cells prevents its localization to centromeres and results in elevated Ctf3 sumoylation.

In principle, elevated Ctf3 sumoylation in *ctf3-2A* cells could reflect disrupted Ulp2 activity on soluble Ctf3 that is not incorporated into centromeric kinetochore particles. To test this possibility, we extended the sumoylation analysis to Mcm21, a member of the four-protein inner kinetochore COMA complex that does not depend on the Ctf3c for its centromeric localization (Measday et al., 2002). Sumoylated Mcm21 was observable in SUMO-enriched wild type cell extracts, and the sumoylation levels, especially levels of poly-sumoylated Mcm21, were more prominent in the *ctf3-2A* background (Figure 3D). The *ulp2-SIM-3A, ulp2-KIM-3A*, and *ulp2-SIM-3A-KIM-3A* double mutant all produced pronounced Mcm21 hyper-sumoylation in otherwise wild type cells. Consistent with the pulldown assay described in Figure 3A, the appearance of sumoylated Mcm21 was elevated further when these Ulp2 mutations were combined with *ctf3-2A*. This was especially true for the *ulp2-SIM-3A* mutant, in agreement with the notion that maximal Ulp2 recruitment to the inner kinetochore depends on cooperative Ulp2 binding to both Ctf3c and sumoylated kinetochore proteins.

### The ctf3-2A protein is specifically deficient in centromeric Ulp2 recruitment

To determine whether the localization defect in *ctf3-2A* cells is restricted to Ulp2, we investigated kinetochore assembly. Cells lacking Mcm16, a Ctf3c component required for its assembly and function, display elevated Mcm21 sumoylation (Suhandynata et al., 2019). The phenotype is similar in degree to that seen in *ctf3-2A*. To verify that the Ctf3-2A protein supports Ctf3c formation and kinetochore recruitment, we imaged cells expressing Mcm22-GFP (Figure 4A). Wild type cells showed Mcm22-GFP localization identical to what we have described for Ctf3-GFP (Hinshaw and Harrison, 2020). The signal in the *ctf3-2A* cells was indistinguishable from that observed in wild type cells, indicating that the Ctf3-2A protein does not disrupt Ctf3c assembly or kinetochore recruitment. To verify this finding, we performed ChIP-qPCR to assess Ctf3 or Ctf3-2A localization (Figure 4B). Analysis of two centromeres (*CEN3* and *CEN7*) and a third site on the arm of chromosome VIII (*CUP1*) showed no statistically discernable defect in Ctf3-2A localization to *CEN3* and a very small but statistically significant deficit in Ctf3-2A localization to *CEN7*.

**Figure 4.**
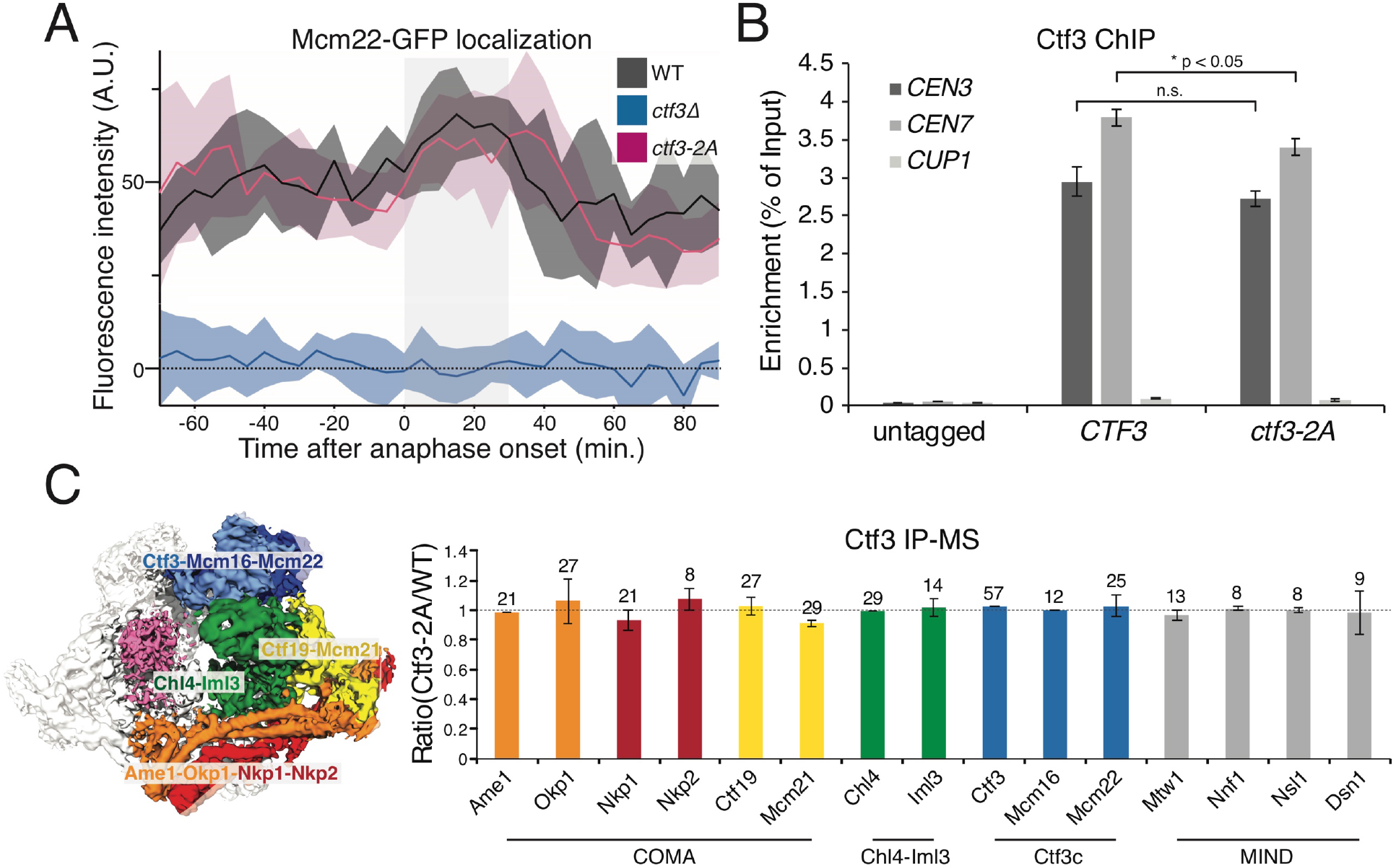
The ctf3-2A mutation does not disrupt kinetochore assembly. A) Mcm22-GFP localization is not perturbed in *ctf3-2A* cells. Kinetochore-associated Mcm22-GFP signal was quantified in asynchronously dividing cells of the indicated genotypes. Mean intensity values from measurements aligned according to the onset of anaphase are plotted as dark lines, and shaded areas indicate 95% confidence intervals. The light gray background demarcates the approximate duration of anaphase as determined by spindle pole body movements (SPB, marked by Spc110-mCherry foci). B) Ctf3 localization is not perturbed in *ctf3-2A* cells. Association of Ctf3 or Ctf3-2A with the indicated genomic loci was assessed by ChIP-qPCR in asynchronously dividing cells (n.s. – not significant; * – *p* ≤ 0.05; Student’s t-test, two tails, equal variance). C) Quantitative mass spectrometry analysis of Ctf3-associated proteins shows that *ctf3-2A* has no detectable effect on the assembly of the Ctf19 complex. The relative abundance of each kinetochore subunit in the purified WT and *ctf3-2A* samples was quantified. Abundance ratios were calculated based on the sum of the intensities of peptides assigned to each protein. Error bars show standard deviations of the top three most abundant peptides, and the number above indicates the number of unique peptides detected for each protein.

We also compared overall kinetochore assembly in *CTF3* and *ctf3-2A* backgrounds. To do so, we purified Ctf3 or Ctf3-2A and associated proteins from exponentially growing cultures and subjected the immunopurified material to quantitative mass spectrometry (Figure 4C). Consistent with the imaging and ChIP-qPCR experiments described above, there was no difference in the levels of kinetochore proteins recovered from extracts from either background. Therefore, under favorable growth conditions, partially elevated inner kinetochore sumoylation in the *ctf3-2A* mutant does not cause kinetochore disassembly.

As two further tests of the impact of the *ctf3-2A* allele on overall kinetochore assembly and its timing, we quantified the kinetochore recruitment of Sgo1-GFP and Scc2-GFP in dividing cells. Sgo1 dissociates from centromeres upon sister kinetochore biorientation in a process that depends on SUMO signaling (Nerusheva et al., 2014; Su et al., 2019). We did not observe a defect in the pattern of Sgo1-GFP localization in the *ctf3-2A*-expressing cells (Figure S3A), indicating that inner kinetochore SUMO regulation is distinct from the pathway that regulates Sgo1 localization. Additionally, although the Ctf19c proteins Chl4 and Iml3 are required for pericentromeric Sgo1 localization in meiosis (Kiburz et al., 2005), we did not observe defective Sgo1 localization in mitotic *ctf19Δ* cells. Scc2 loads cohesin onto chromosomes, and its transient recruitment to kinetochores depends on Ctf3-mediated Dbf4-dependent kinase (DDK) localization in late G1 (Hinshaw et al., 2017). Scc2-GFP localization was unperturbed in *ctf3-2A* mutant cells (Figure S3B). Therefore, elevated kinetochore sumoylation does not impair centromeric cohesin loading or the kinetochore processes required to support this activity, consistent with previous observations in *ulp2Δ* (Bachant et al., 2002).

### Defective Ulp2 localization disrupts chromosome segregation

While inner kinetochore assembly appeared normal in unstressed *ctf3-2A* mutant cells (described above), it remained possible that Ulp2 safeguards kinetochore assembly under stressful conditions, analogous to the essential activity of its human homolog, SENP6. Indeed, genetic complementation experiments showed that Ulp2 mutations found to be important for Ctf3 interaction in biochemical experiments described above (Figure 2) produced a viability defect when combined with the Ulp2-SIM-3A mutation (Figure S4A). To test whether this defect could be partly explained by defective Ulp2 kinetochore localization, we measured the viability of cells exposed to mild DNA replication stress. Although the effect is modest, the *ctf3-2A* mutation sensitizes cells to hydroxyurea (Figure 5A). The effect is pronounced for *ctf3-2A ulp2-SIM-3A* double mutant cells, in which both mechanisms that support Ulp2 kinetochore recruitment have been disabled. Therefore, maximal cellular fitness requires efficient desumoylation of inner kinetochore proteins by Ulp2, and this is especially true when cells are exposed to replication stress.

**Figure 5.**
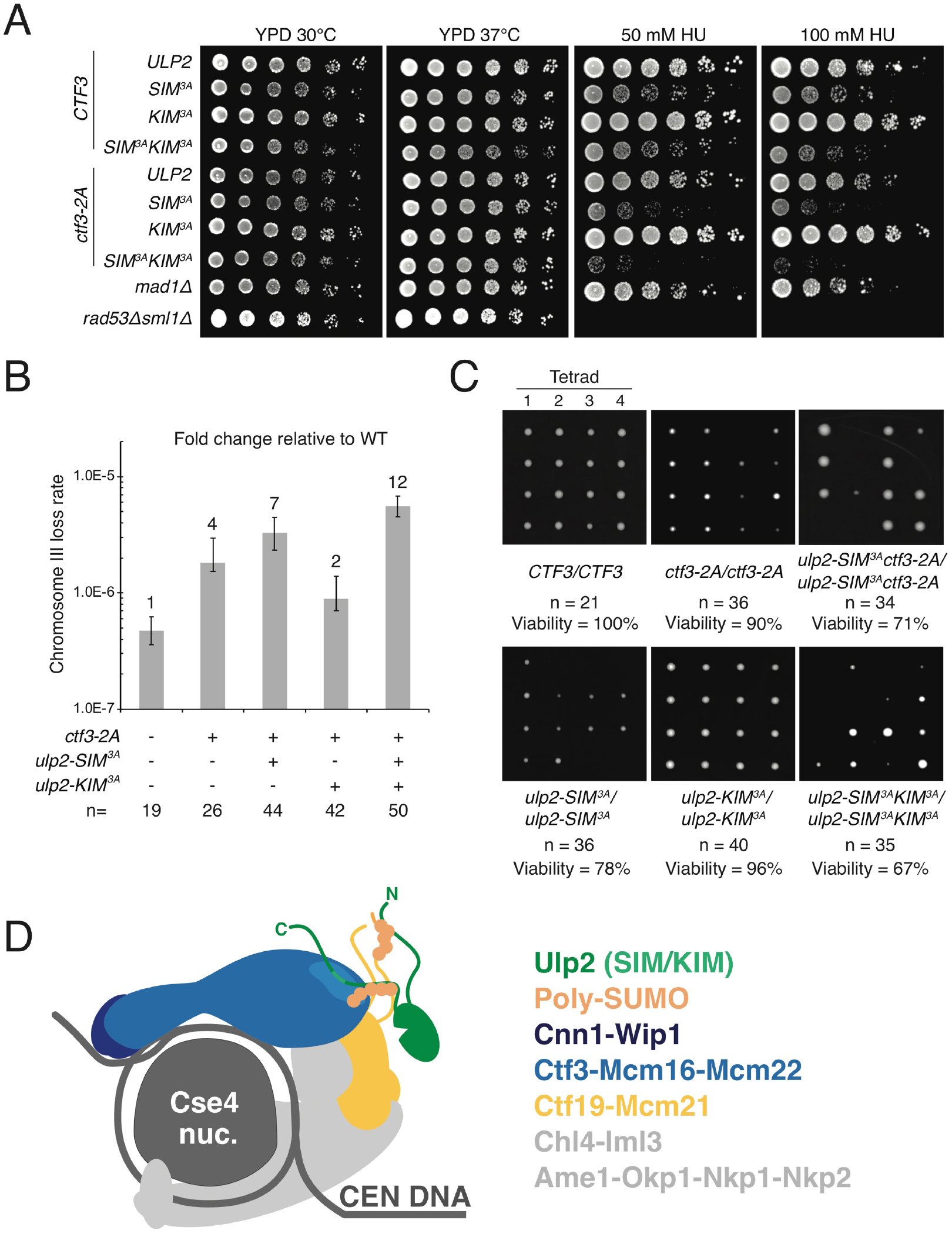
Ulp2 kinetochore activity is required for accurate chromosome segregation. A) Disrupted Ulp2 kinetochore localization in *ulp2-KIM-3A* and *ctf3-2A* cells causes elevated sensitivity to hydroxyurea (HU), and this sensitivity becomes more pronounced with the additional *ulp2-SIM-3A* mutation. 5-fold dilution series are shown. B) Mutations affecting Ulp2 localization cause elevated rates of chromosome III loss. Rates were measured using multiple independent cultures for each mutant. Loss rates are plotted, and numbers above bars show fold changes relative to wild type cells. Error bars show 95% confidence intervals for the median of loss rate. C) Representative images of the products of tetrad dissections of the indicated homozygous diploid parental strains. Spore viability is the ratio between the number of viable spores and the total expected spores. n indicates the number of tetrads scored for each genotype. D) Cartoon model showing Ulp2 association with the inner kinetochore. The image is adapted from Hinshaw and Harrison, 2020 (Cse4 nuc. – Cse4 nucleosome; CEN DNA – centromeric DNA). Extended N- and C-terminal peptides from Ctf19, Mcm21, and Ulp2 are shown as colored lines. Ulp2-N and −C are labeled, and the Ulp2 catalytic domain is shown as a green oval.

The most likely explanation for impaired fitness in *ctf3-2A ulp2-SIM-3A* mutant cells is impaired chromosome segregation (Ryu et al., 2016; Suhandynata et al., 2019). To test this idea, we measured chromosome segregation using a mating type faker assay and found that *ctf3-2A* cells missegregate chromosome III more frequently than wild type cells (Figure 5B). The effect was similar in *ulp2-SIM-3A* cells and more pronounced in a strain lacking all three kinetochore interaction motifs (*ctf3-2A ulp2-SIM-3A-KIM-3A*). Meiotic chromosome segregation is especially sensitive to defective kinetochore regulation. We tested spore viability in homozygous wild type, *ctf3-2A, ulp2-SIM-3A, ulp2-KIM-3A, ulp2-SIM-3A ctf3-2A*, and *ulp2-SIM-3A-KIM-3A* mutant cells (Figure 5C). All mutations increased the frequency of inviable spores. Cells carrying the *ulp2-SIM-3A* mutation had a pronounced defect, consistent with SIM-dependent homeostatic control of both kinetochore and nucleolar sumoylation (de Albuquerque et al., 2018). The mild but observable sporulation defect seen in *ctf3-2A* mutant cells and the more pronounced defect in the *ulp2-KIM-3A ctf3-2A* double mutant cells indicates a role for regulated kinetochore desumoylation in normal meiotic chromosome segregation.

## DISCUSSION

Ulp2 regulates multiple nuclear processes. Interactions between subnuclear structures and the enzyme itself determine specificity and timing (Liang et al., 2017; Srikumar et al., 2013; Suhandynata et al., 2019). Efficient kinetochore recruitment depends on two conserved elements in the non-catalytic C-terminal tail of Ulp2 (Suhandynata et al., 2019). One of these peptides binds SUMO chains, and the other binds the Ctf3c. We have identified the surface of the Ctf3 protein responsible for the latter interaction by solving the cryo-EM structure of the complex. The Ctf3 surface is conserved, indicating the interaction may be ancient, and mutations at the surface perturb the Ctf3c-Ulp2 interaction both in reconstituted biochemical systems and in cells.

The Ctf3-Ulp2 interaction requires both electrostatic and nonpolar contacts. The former depends on conserved Ulp2-KIM aspartate side chains, while the latter depends on conserved hydrophobic Ulp2-KIM side chains. These contacts are distinct from those that mediate SIM-SUMO contacts, despite the fact that the Ulp2-KIM and −SIM peptides have similar primary sequences. Indeed, the Ulp2-KIM has no detectable SUMO-binding activity (de Albuquerque et al., 2018), and the Ulp2-SIM does not bind the Ctf3c. Conventional SIM peptides interact with SUMO-1 by beta sheet extension (Namanja et al., 2012). The Ctf3c site we have identified precludes this mode of interaction. Thus, the Ulp2-KIM and Ulp2-SIM are mechanistically and structurally distinct recruitment motifs that, together, enable fine-tuning of Ulp2 kinetochore activity in response to cellular and environmental perturbations.

The strength and timing of Ulp2 activity at the inner kinetochore varies during the cell cycle (Suhandynata et al., 2019). Ulp2 recruitment, as we have shown here and previously, depends on distinct KIM-Ctf3c and SIM-SUMO interactions. Only the KIM-Ctf3 interaction is specific to the inner kinetochore. The two-site mechanism implies homeostatic regulation: SUMO chain cleavage disfavors Ulp2 recruitment. Steady-state inner kinetochore sumoylation is therefore set by a competition between Ulp2 and fluctuating activities of nuclear SUMO ligase complexes. Kinetochore kinases may also contribute to observed cell cycle-dependent changes in Ulp2 activity (Suhandynata et al., 2019). The extended regions of Ctf19 and Mcm21, which are located near the Ctf3 conserved surface in the assembled Ctf19c, are both phosphorylated and are therefore good candidate mediators of this regulation (Albuquerque et al., 2008; Hinshaw et al., 2017). Ulp2 itself is also phosphorylated at sites near the C-terminal SIM and KIM peptides described here (Baldwin et al., 2009). The precise functions of most of these modifications have not been described, and their cell cycle-dependent activities are promising topics for further study.

SUMO metabolism regulates interlocking nuclear processes. Alleles that separate these overlapping functions therefore constitute important tools. In addition to influencing chromosome segregation by targeting inner kinetochore proteins, Ulp2 regulates essential nuclear processes, including DNA replication, nucleolar gene silencing, and recovery after DNA damage-induced arrest (de Albuquerque et al., 2016; Liang et al., 2017; Schwartz et al., 2007; Wei and Zhao, 2016). Its known substrates in these processes include the Mini-Chromosome Maintenance (MCM) complex, the nucleolar RENT complex, the DNA topoisomerase Top2, and the DDK subunit Dbf4 (Bachant et al., 2002; de Albuquerque et al., 2016; Liang et al., 2017; Psakhye et al., 2019; Wei and Zhao, 2016). The structure we have described enables precise manipulation of Ulp2 activity at the inner kinetochore without perturbation of these related functions.

We have used the new *CTF3* alleles to investigate the influence of sumoylation on kinetochore assembly and chromosome segregation. In particular, unperturbed Scc2 localization indicates the *ctf3-2A* cells maintain DDK recruitment to the kinetochore in late G1/early S phase. Likewise, distinct recruitment mechanisms probably control Ulp2 activity in the pathways enumerated above. Independent regulation enables variable tuning of Ulp2 activity appropriate for each context and provides an evolutionary explanation for extended N- and C-terminal regulatory segments that flank a conserved enzymatic core.

Both Ulp2 (*SMT4*) and SUMO itself (*SMT3*) were originally identified as high copy genetic suppressors of a temperature-sensitive *MIF2* allele, an essential component of the inner kinetochore (Meluh and Koshland, 1995). The human Ulp2 homolog, SENP6, removes SUMO modifications from CENP-I and other inner kinetochore proteins (Fu et al., 2019; Liebelt et al., 2019; Mitra et al., 2020; Wagner et al., 2019), and this activity is required for kinetochore stability (Mukhopadhyay et al., 2010). Identification of a conserved kinetochore site that mediates Ulp2 recruitment and directs its activity to a specific subset of inner kinetochore substrates provides a possible explanation for these findings. A similar pathway may function in multicellular organisms to ensure kinetochore maintenance during extended periods of cellular quiescence or arrested growth.

## ACKNOWLEDGMENTS

We thank the staff at the Harvard Cryo-Electron Microscopy Center for Structural Biology for help with high-resolution cryo-EM data collection, particularly Richard Walsh and Sarah Sterling. We thank Shaun Rawson for real-time movie processing. We thank SBGrid for computational support. We thank Jennifer Waters and the staff at the Nikon Imaging Center at Harvard Medical School for light microscopy support. This work was supported by NIH GM116897, OD023498, and University of California CRCC faculty seed grant to H.Z. S.M.H. is an HHMI fellow of the Helen Hay Whitney Foundation. S.C.H. is an Investigator of the Howard Hughes Medical Institute.

## DECLARATION OF INTERESTES

The authors declare no competing interests.

## DATA AVAILABILITY

The accession numbers for the cryo-EM structure are EMD-23216 (map) and PDB 7L7Q (model) and have been deposited in the EMDB and PDB, respectively.

---

**Figure S1.**
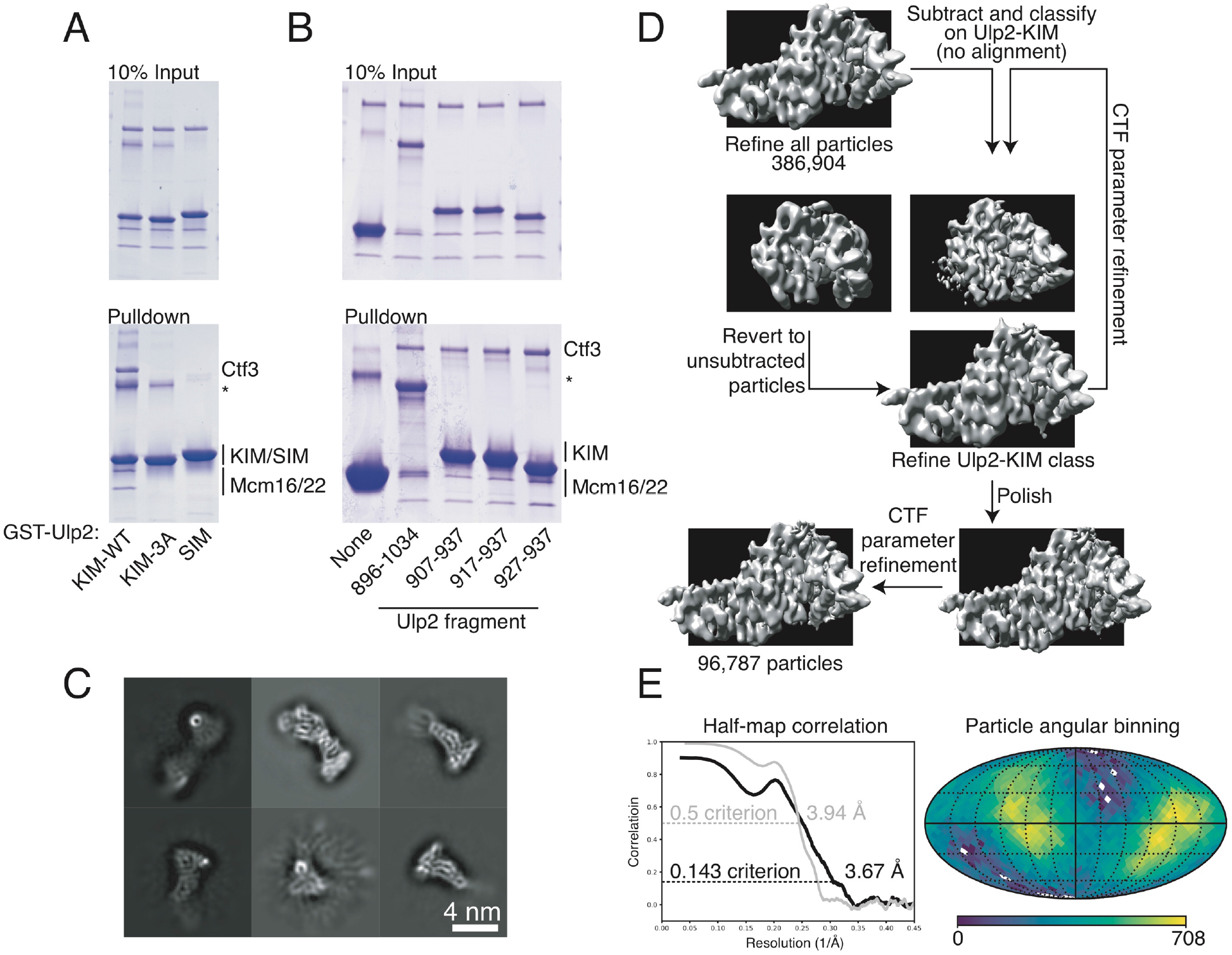
Identification of a minimal Ulp2-KIM and Ctf3c-Ulp2 structure determination. A) The Ctf3c-Ulp2 interaction is specific. GST pulldown was used to detect the interaction between the indicated GST-Ulp2 fusion proteins (KIM-WT – Ulp2-927-937; KIM-3A – Ulp2-927-937 with 3A substitutions; SIM – Ulp2-717-734) and the Ctf3c (* – contaminant in GST fusion protein preparations). B) An 11-residue Ulp2-KIM peptide binds the Ctf3c. GST pulldown was used to detect the interaction between the indicated GST-Ulp2 peptides and the Ctf3c (* – contaminant in GST fusion protein preparations). C) Two-dimensional class averages from initial stages of particle processing. D) CryoEM data processing for the Ctf3c-Ulp2-KIM complex. The model shown as the starting point here is the product of three-dimensional refinement after merging of the two datasets (Table S1). Prior to merging, two- and three-dimensional classification steps were used to select particles contributing to the Ctf3c module shown. Unsharpened volumes are shown. E) Fourier shell correlation (black – half-map to half-map; gray – model to map) and Euler angle distribution for contributing particle images are shown. Angular binning density is presented on a logarithmic scale.

**Figure S2.**
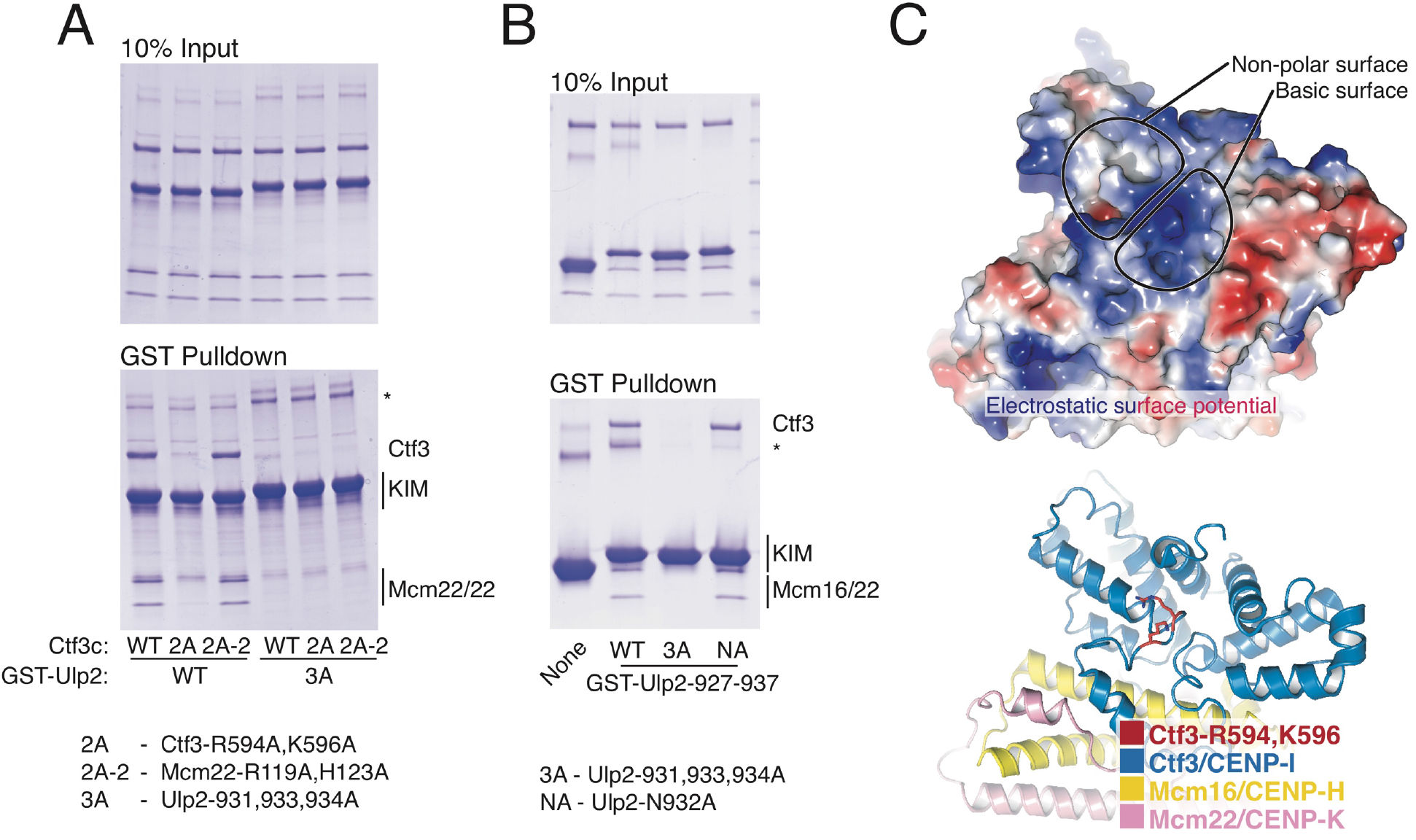
Determinants of the Ctf3c-Ulp2 interaction. A) Biochemical identification of the *ctf3-2A* mutation. GST pulldown was used to detect binding between the indicated Ctf3c samples (WT – wild type; 2A – Ctf3-R594A,K596A; 2A-2 – Mcm22-R119A,H123A; * contaminants). The 2A-2 mutant Ctf3c probes a second Ctf3c surface that does not bind Ulp2. B) Ulp2-N932 is not required for Ctf3c interaction. GST pulldown with the indicated GST-Ulp2-927-937 peptides (WT – wild type; 3A – Ulp2-V931A,L933A,I934A; NA – Ulp2-N932A) was used to probe Ctf3c binding. C) Two views of the Ctf3c showing the Ulp2 binding site. Top panel shows the Ctf3c surface colored according to electrostatic surface potential (blue – positive; red – negative; gray – nonpolar). Bottom panel shows a cartoon view of the Ctf3c in the same orientation with the amino acid residues substituted by alanine in the Ctf3-2A protein colored red.

**Figure S3.**
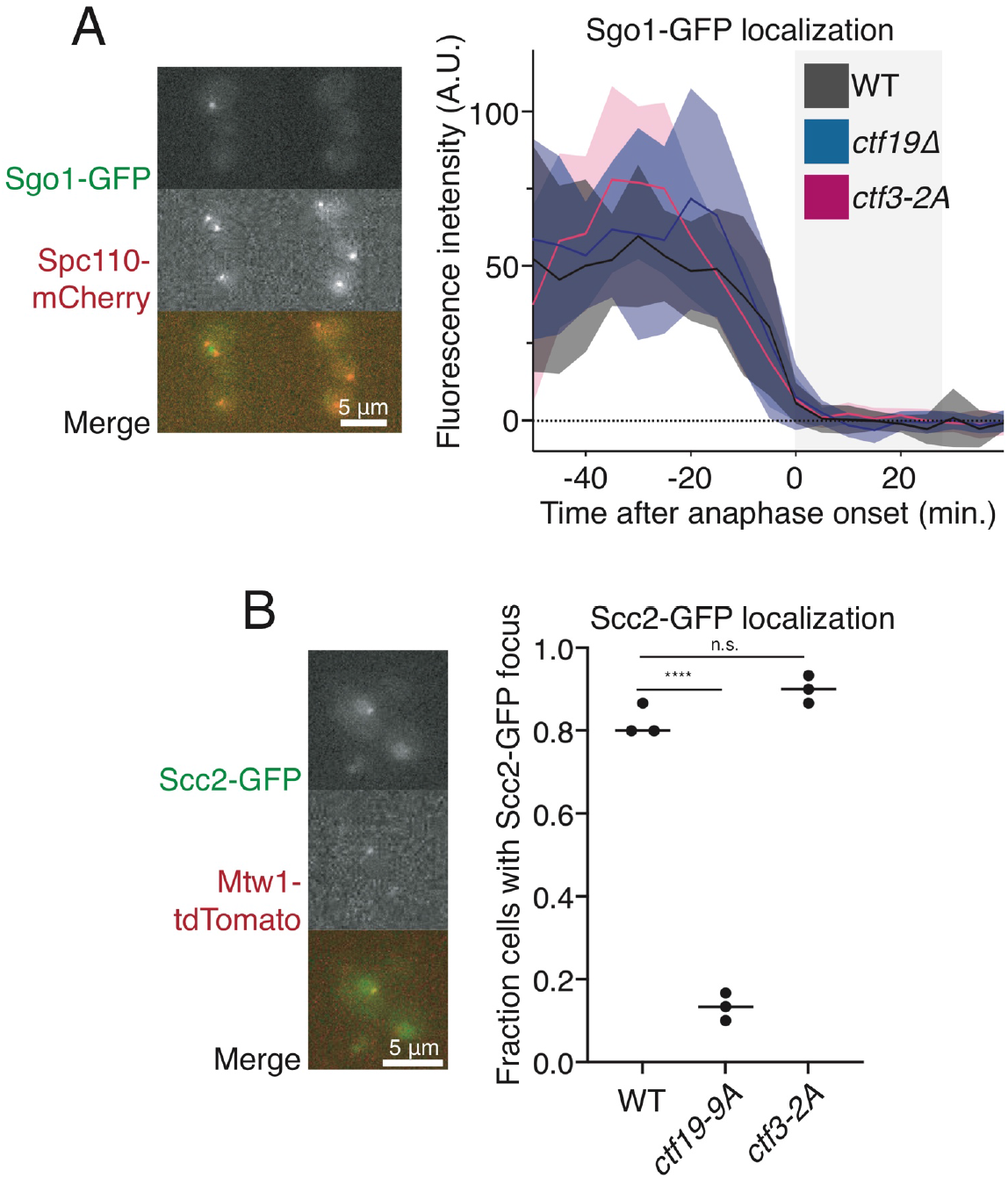
Consequences of kinetochore assembly in wild type and ctf3-2A cells. A) The *ctf3-2A* mutation does not perturb Sgo1 localization or its dissociation from centromeres upon anaphase initiation. Sgo1-GFP inter-kinetochore intensity was measured as for Mcm22-GFP (Figure 4A). Solid lines show mean intensity values, and shaded areas show 95% confidence intervals. The light gray background demarcates the approximate duration of anaphase as determined by SPB movements. Sgo1-GFP localizes to a punctate structure between Spc110-mCherry SPB markers, and the signal dissipates when SPBs separate on the anaphase spindle. A representative dividing cell is shown before (left) and during (right) anaphase. B) The *ctf3-2A* mutation does not perturb Scc2 localization. The presence or absence of an Scc2-GFP focus was quantified in at least 30 dividing cells for three distinct experiments for each indicated genotype (n.s. – not significant; **** – p ≤ 0.0001; Student’s t-test, two tails, unequal variance). *ctf19-9A* disrupts Scc2 kinetochore localization (Hinshaw et al., 2017). Scc2-GFP appears as a transient kinetochore-associated focus as cells enter S phase. A representative cell is shown.

**Figure S4.**
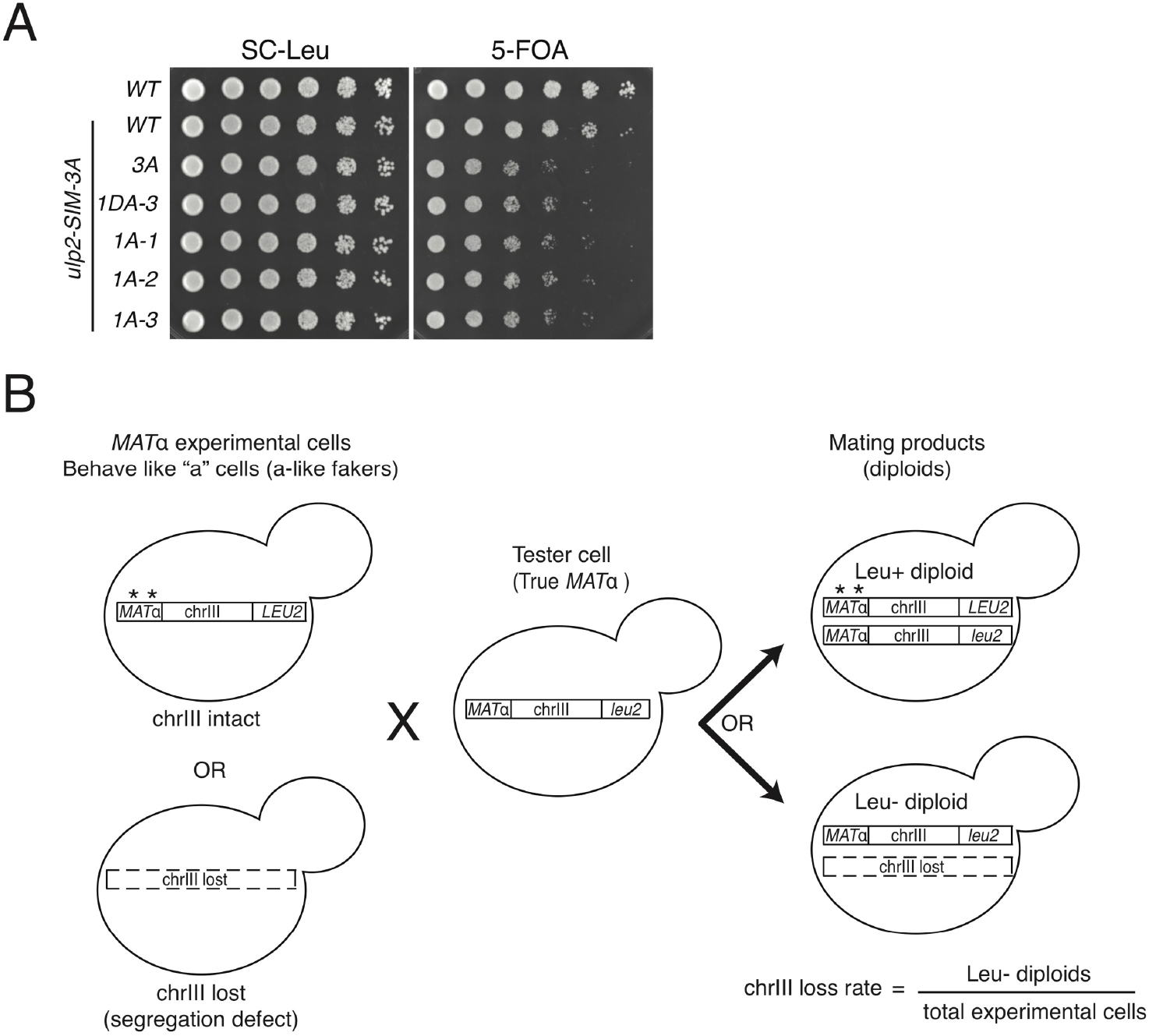
Viability of Ulp2 mutant strains and chromosome segregation assay. A) Growth of various *ulp2* mutants. Loss of a complementing *ULP2* plasmid was induced with 5-fluoroorotic acid (5-FOA). In addition to the indicated *ulp2* mutations, the experimental strains carried the *ulp2-SIM-3A* mutation on the same experimental allele. B) Schematic showing the measurement of chromosome loss rate using the quantitative mating assay (Figure 5B). Mat alpha experimental cells that mutated in MATα locus or lost whole chromosome III behave like “a” cells and are able to mate with Mat alpha tester cells to form diploids. Diploid cells generated by chromosome III loss are Leu negative due to the coincident loss of *LEU2* on the opposite arm of chromosome III to *MATα*, which distinguishes mating events from mutations that result in Leu positive colonies.

## METHODS

### Yeast strain construction and growth conditions

Yeast strains were generated by standard methods (Table S2) and grown in rich medium with additives or dropout medium (synthetic complete (SC), Sunrise Science Products) at 30 °C unless otherwise indicated. Mutant strains were generated by lithium acetate-mediated integration of PCR products (Longtine et al., 1998). Mutations were verified by Sanger sequencing (new alleles) or PCR genotyping (deletion strains). Standard molecular biology techniques were used to generate plasmids (Table S3), and details for their construction are available upon request.

### Recombinant protein production

Recombinant Ctf3c protein samples were purified from *E. coli* Rosetta 2(DE3)pLysS cells as described previously (Hinshaw and Harrison, 2019). N-terminal His6 tags were removed from all Ctf3c samples by incubation with TEV protease at room-temperature for 3 hours after ion exchange chromatography as described (Hinshaw and Harrison, 2019). Plasmids used for Ctf3c expression contain a single mRNA encoding the three subunits, each controlled by translation start and stop sequences. Ctf3 mutations were created by two-step overlapping PCR amplification of the polycistronic insert before reinsertion into the original destination vector by ligation-independent cloning. Recombinant Cnn1-Wip1 protein sample used for cryo-EM was the same as used previously (Hinshaw and Harrison, 2020).

GST-tagged Ulp2 fragments were expressed in *E. coli* Rosetta 2(DE3)pLysS cells (Millipore) grown to an optical density of ~0.8 at 37 °C before shifting the temperature to 18 °C and induction of protein expression with 0.4 mM IPTG. After ~16 hours, cells were harvested, resuspended in buffer D800 (20 mM HEPES, pH 7.5; 10 mM imidazole, pH 8.0; 800 mM NaCl; 2 mM β-mercaptoethanol; 10% glycerol) with protease inhibitors (2 μg/ml each aprotinin, leupeptin, and pepstatin; 1 mM each PMSF and benzamidine) and frozen at −80 °C. His6-GST fusion proteins were purified by metal affinity chromatography as for Ctf3c samples except that protease inhibitors (0.5 μg/ml each aprotinin, leupeptin, and pepstatin) were included in the metal chromatography elution buffer. The resulting eluate was concentrated by ultrafiltration and applied to a Superdex 200 column (10/300 GL, GE) equilibrated in GF100 buffer (20 mM Tris-HCl, pH 8.5, 100 mM NaCl, 1 mM DTT). Peak fractions were pooled, concentrated by ultrafiltration, and stored in 5% glycerol at −80 °C until use.

### GST pulldown assay

The indicated proteins were mixed and incubated on ice for one hour in a total volume of 20 μL GST pulldown buffer (20 mM tris pH 8.5; 150 mM NaCl; 1 mM TCEP; 0.05% NP-40 substitute). After incubation on ice, 10 μL glutathione-sepharose beads were added, and the samples were mixed end-over-end for one hour at 4 °C. Beads were washed four times with pulldown buffer, and the bound material was eluted by boiling with SDS-PAGE buffer.

### Cryo-EM sample preparation

Purified Ctf3c (120 picomoles) was incubated with an equimolar amount of Cnn1-Wip1 and a 3-fold molar excess of the FITC-Ulp2-KIM peptide (fITC-AHA-DDSDVNLIGSS; Tufts University Core Facility). After incubation on ice for one hour, the mixture was subjected to gel filtration on a Superdex 200 column (S200 increase 5/150 GL; GE) equilibrated with GF150-MALS (20 mM Tris pH 8.5; 150 mM NaCl; 1 mM TCEP; 1 mM EDTA; 0.05% sodium azide). Absorbance at 488 nm confirmed the presence of the FITC-Ulp2 peptide. Equivalent gel filtration experiments using a biotin-labeled Ulp2-KIM peptide coupled with analysis of the eluted material by dot blot using streptavidin-HRP further confirmed the retention of the peptide during gel filtration. For all gel filtration experiments, 50 μL fractions were collected manually and analyzed by SDS-PAGE.

Protein concentration after gel filtration was measured by absorbance at 280 nm, and a fraction containing ~0.5 μg/ml protein was selected for analysis by cryo-EM. 3.5 μL of the eluate was applied to a cryo-EM grid (C-flat 2/1, glow discharged for 30 sec. at 15 mA). The grid was plunged into liquid ethane after blotting from both sides for 4 sec. using a Cryoplunge 3 instrument (Gatan). Vitrified samples were screened for ice thickness and sample distribution using an F20 electron microscope (FEI) operating at 200 kV.

### Cryo-EM data collection and structure determination

Grids were mounted and imaged as described previously (Hinshaw et al., 2019) on a Krios G3i (Gatan) instrument operating at 300 kV controlled by SerialEM (Mastronarde, 2005). The sample was illuminated with a 1 μm beam in nanoprobe mode with a pre-camera energy filter (Gatan) slit width of 20 eV. Movies were collected on a K3 detector (Gatan) in counting mode with a pixel size of 0.825 μm. 48 frames were collected per 2.4 second movie, with a total dose of 52 electrons/Å^2^ divided equally among the frames. The defocus range was set between −1.2 and −3.0 μm. Using image shift and real-time coma-correction by beam tilt as implemented in SerialEM, either nine or four holes were visited at each stage position, and five movies were taken per hole, giving a total of 45 or 20 movies per stage movement.

Initial image processing steps were carried out using an in-line pipeline that uses the RELION 3.0 job scheduling function (Zivanov et al., 2018). Movies were aligned using Motioncor2, and CTF parameters were estimated using CTFFIND4 (Rohou and Grigorieff, 2015; Zheng et al., 2017). All subsequent steps were carried out in RELION 3.1. Initial particle coordinates were determined using 2-dimensional class averages showing the Ctf3c-Cnn1-Wip1 complex as references. Extracted and binned particles were subjected to multiple rounds of 2-dimensional classification to select a subset showing features corresponding to the Ctf3c. Whereas we previously included only 2-dimensional class averages showing the Ctf3-N module (Hinshaw and Harrison, 2020), which contains Cnn1-Wip1, we did not apply this strict filter for the reconstructions presented here. In the course of selecting 2-dimensional class averages suitable for further analysis, we noticed the frequent appearance of a bright punctate signal coincident with the C-terminal end of Ctf3. Because we observed a similar signal corresponding to the location of the extended Cnn1-N peptide in initial class averages that contain this module, we presume this signal corresponds to a peptide contacting the air-water interface. The position of the signal matches the projected position of the Ulp2-KIM peptide. Exclusion of class averages in which this punctate signal was dominant and therefore interfered with intensity normalization and particle alignment was essential for the eventual determination of high-quality three-dimensional density maps.

Particles contributing to favorable 2-dimensional class averages were subjected to 3-dimensional classification using an initial model derived from the known structure of the Ctf3c lacking the Ctf3 N-terminal module, Cnn1, and Wip1 (Hinshaw et al., 2019). The resulting particles were then subjected to iterative rounds of 3-dimensional refinement, CTF and optical parameter fitting, and alignment-free classifications. The Ulp2-KIM peptide appeared as additional density not observed in the published high-resolution Ctf3c-Cnn1-Wip1 structure. The extra density is adjacent to the C-terminal end of Ctf3. We used iterative rounds of signal subtraction and alignment-free classification to select particles contributing meaningfully to this extra density.

The resulting density map is nearly identical to those reported previously for the corresponding part of the Ctf3c. Additional density near the Ctf3 C-terminus is attributable to the Ulp2-KIM. We truncated the Ctf3c model, determined at higher resolution, and subjected the modified coordinates to a single round of refinement against the unsharpened final density map (Phenix v1.17.1 Real Space Refine) (Afonine et al., 2018). The starting and final models are highly concordant (RMSD 0.955), indicating Ulp2-KIM binding does not induce a conformational change in the Ctf3c. Although the density for the Ulp2-KIM was visible throughout refinement and particle selection, it does not permit unambiguous placement of Ulp2-KIM amino acid side chains. We therefore opted to omit an explicit Ulp2-KIM model from our coordinates and figures.

### Fluorescence polarization assay

The indicated purified Ctf3c samples were mixed with the FITC-Ulp2-KIM peptide (Ulp2-927-937). The peptide was reconstituted in gel filtration buffer with 150 mM NaCl and stored protected from light at −20 °C until use. The final peptide concentration used for binding assays was 40 nM. Binding reactions were prepared in a black 384-well plate in gel filtration buffer with 150 mM NaCl (20 mM Tris, pH 8.5; 150 mM NaCl; 1 mM TCEP) and incubated at room temperature in the dark for 15 min. before reading in an M5e Spectramax Plus plate reader (Molecular Devices) in fluorescence polarization mode (Ex. 485 nm, Em. 590 nm, 3 readings per well) at room temperature. Three independent reactions were prepared for each measurement, and averages are shown. The experiments shown were carried out at least twice with equivalent results. Recombinant GST-Ulp2-927-937 proteins used for competition experiments were added at a final concentration of 5 μM to the binding reactions at the indicated Ctf3c concentrations. A single specific binding event was modeled after linear correction of all measurements for polarization signal in the absence of Ctf3c.

### Sequence and conservation analysis

Sequence alignments were constructed using MAFFT and displayed with JalView (Katoh et al., 2019; Waterhouse et al., 2009). Inputs for Ctf3 multiple sequence alignments were adapted from van Hoof (van Hooff et al., 2017). Conservation was mapped to the Ctf3 protein structure using the ConSurf server (Ashkenazy et al., 2016). The Ulp2-KIM sequence logo was generated using the WebLogo 2.8.2 server (Crooks et al., 2004).

### Pulldown analysis of kinetochore sumoylation

The dUlp1 affinity method, described previously (Suhandynata et al., 2019), was used to analyze sumoylation of Mcm21 and Ctf3 in various mutant strains. Briefly, each yeast strain was grown in 100 mL YPD medium to an OD_600_ of 0.8 and harvested. Cell pellets were washed with PBS buffer, supplemented with protease inhibitors (2 mM phenylmethylsulfonyl fluoride, 200 μM benzamidine, 0.5 μg/mL leupeptin, 1 μg/mL pepstatin A), 20 mM N-Ethylmaleimide and 20 mM Iodoacetamide. Cells were then broken via glass bead-beating, and the clarified cell extracts were collected following centrifugation at 14,000 rpm for 30 min. To purify total sumoylated proteins, each clarified cell extract (0.8 mL at a protein concentration of ~12 mg/mL) was incubated with 20 μL dUlp1 resin for 2 hours at 4°C with gentle rotation. The dUlp1 resin was then washed six times with 1mL PBS containing 0.2% NP-40. Bound proteins were eluted by boiling in 25 μL of 2×LDS sample buffer (NuPAGE LDS Sample Buffer, Invitrogen). Anti-Protein A Western blot was performed to detect sumoylated Mcm21 or Ctf3 species in the purified material, which show slower electrophoretic mobility due to their increased molecular weight relative to the corresponding un-sumoylated proteins in the input samples.

### Purification of Ctf3-associated proteins and quantitative mass spectrometry analysis

Quantitative mass spectrometry was used to compare Ctf3-associated proteins purified from WT and *ctf3-2A* mutant. Briefly, cells expressing wild-type Ctf3-TAF were grown in 1 liter of heavy Lys/Arg containing synthetic media, while the *ctf3-2A-TAF* mutant was grown in synthetic media containing light Lys/Arg till OD_600_ ~ 0.5 and then these cells were harvested and handled separately. Cells were broken via glass beads beating in the PBS with 0.2% NP-40, supplemented with protease inhibitors, and the clarified cell extract was collected following centrifugation as described above. Equal amounts of cell extracts (~ 100 mg total proteins each) were used to purify Ctf3-associated proteins as follows. First, each clarified cell extract was incubated with 150 μL of human IgG Sepharose beads (Cytiva) at 4°C for 3 hours. The beads were washed four times with 1 mL lysis buffer, and then incubated with 3 μg TEV protease for overnight at 4°C. The next day, the TEV-eluted samples were collected and incubated with 20 μL of Anti-Flag agarose beads (Sigma) for 2 hours at 4°C. The beads were washed four times by ice-cold lysis buffer and bound proteins were eluted with 0.1 M Glycine-HCl (pH ~1-2). The eluted materials from both Ctf3 purifications were neutralized and combined as one. To the combined sample, 10 mM DTT was added to reduce disulfide bonds in proteins, which were then alkylated with 30 mM iodoacetamide. To digest the proteins, 1μg trypsin (Promega) was added to the sample for overnight incubation at 37 °C. The digested peptides were then processed for analysis using a Thermo Scientific Orbitrap Fusion LUMOS Tribrid mass spectrometer. Methods used for mass spectrometry analysis are described elsewhere (Suhandynata et al., 2019). To quantify Ctf3-associated kinetochore proteins, several additional data filtering criteria were applied, including: 1) a cutoff score for peptide identification probability was set to at 0.9; 2) parental ion mass accuracy was < 10 ppm; and 3) the spectral intensity was >1000. In case of redundant identification of a peptide, the peptide with the most intense signal was kept for quantification. Each protein was quantified with ≥ 3 unique peptides, and the sum of the abundances of these peptides was taken to determine the protein abundance ratio between Ctf3-WT and *ctf3-2A*. Standard deviation was calculated using the abundance ratios of the top three most abundant and unique peptides of each protein. Complete list of Ctf3-associated proteins and their abundance changes are in Table S4.

### Fluorescence microscopy and analysis

Fluorescence microscopy experiments were carried out as described previously (Hinshaw and Harrison, 2020). The indicated strains were grown overnight in synthetic complete medium supplemented with 20 μg/ml additional adenine (SCA). Before imaging, saturated cultures were diluted 1:100 (*v*:*v*) and grown for ~6 hours before immobilization on concanavalin A-coated cover slips, which were subsequently mounted in a Toaki Hit stage-top incubator set to 30 °C at the specimen level with high humidity. Images were collected on a Nikon Ti2 inverted microscope equipped with an Olympus PlanApo 60x/1.42 NA oil immersion objective, the Perfect Focus System, and a Hamamatsu Flash4.0 V2+ sCMOS camera controlled with NIS Elements software (Nikon). Optical paths were: GFP – SpectraX Cyan (illumination), Lumencor 470/24 (excitation), and Semrock FF03 525/50 (emission); mCherry – SpectraX GreenYellow (illumination), Lumencor 525/25 (excitation), and Semrock FF02 641/75 (emission). Nine z-heights (0.4 μm spacing) were taken per stage position. Image stacks were taken at five-minute intervals for at least two hours for each imaging session.

Maximum z-projection images were analyzed as previously described (Hinshaw and Harrison, 2020). Mean intensity within a 6-pixel circle centered on the kinetochore focus was normalized by subtracting a paired measurement from a nearby nuclear position at each timepoint and for each analyzed cell. Individual intensity profiles for dividing cells were aligned according to the onset of anaphase, and average intensity values were plotted using GraphPad Prism v8.4.1 software (GraphPad). All measurements and display items were created using Fiji (Schindelin et al., 2012). For display items, brightness and contrast were adjusted equally for all images corresponding to the same GFP fusion protein, and panels were assembled in Adobe Photoshop software.

### Chromatin immunoprecipitation (ChIP)

To evaluate localization of Ulp2 and Ctf3 to the centromere, ChIP was performed as described previously (Meluh and Koshland, 1997; Suhandynata et al., 2019). Briefly, yeast cultures (150 mL, enough for three immunoprecipitation experiments) were grown to an OD_600_ of 0.8 and cross-linked for 15 min with 1% formaldehyde at room temperature. Whole cell lysates were prepared in 0.8 mL of ChIP lysis buffer (50 mM Hepes pH 7.6, 140 mM NaCl, 1mM EDTA, 1% Triton, 0.1% DOC) by glass beads beating and sonicated to shear the genomic DNA to an average size of 300-500 bp. Immunoprecipitation was performed using 50 μL Dynabeads Protein G (ThermoFisher) and 3 μL anti-Flag antibody M2 (F3165, Sigma). The beads were washed once with 1 mL lysis buffer, twice with 1 mL washing buffer (100 mM Tris-Cl pH 8.0, 250 mM LiCl, 0.5% NP-40, 0.5% deoxycholate, 1 mM EDTA), and once with 1 mL TE buffer (10 mM Tris-Cl pH 8.0, 1 mM EDTA). The input and immunoprecipiated DNA were reverse cross-linked at 60 degrees for 12 hours and then purified using a QIAquick PCR Purification kit (QIAGEN). The input DNA was diluted 1:100, and immuno-purified DNA was diluted 1:10. qPCR was done using SYBR Green 2x master mix (KAPA Biosystems) on a Roche LightCycler 480 system. Three independent immunoprecipitation experiments were performed. qPCR primer sequences are available upon request.

### Cell growth and meiotic spore viability assays

For the spot assay, cells were grown in YPD to OD_600_ ~1 and then diluted to an OD_600_ of 0.2. Five-fold serial dilutions were made using a sterile 96-well plate with sterile water. 5 μL of each dilution was then spotted on either YPD plates or hydroxyurea plates (YPD supplemented with the indicated concentrations of hydroxyurea). For the plasmid shuffling experiment (Figure S4A), cells were grown in SC-Leu media handled as described above. 5 μL of each dilution was spotted on either SC-Leu or 5-FOA plates (SC supplemented with 0.1% 5-FOA). Plates were incubated at 30 °C for two or three days. Plate images were taken using a Bio-Rad ChemiDoc MP imaging system. Tetrad dissections of diploids containing *ulp2* or *ctf3-2A* homozygous mutants were performed using a dissecting microscope (Singer). Spore viability was calculated based on the ratio between the numbers of observed spores and the total expected spores given 100% viability.

### Mitotic chromosome segregation assay

Chromosome loss rate was measured using a previously described quantitative mating assay (Vaezzadeh et al., 2010). As outlined in Figure S4B, *MATα ARG2 LEU2 ura3* experimental haploids were mated to a tester strain (HZY601: *MATα arg2 URA3*) by mixing ~1×10^7^ log-phase cells of each strain on a filter membrane (0.8 μm MCE Membrane Filter, MF-Millipore) on top of a YPD plate for 5 hours at 30 °C. Cells were resuspended in 1 mL of sterile dH2O and collected. Successfully mated diploid cells were selected by plating 10-100% of the cell resuspension onto SC-Arg-Ura plates, such that the number of colonies was between 100 and 200. The colonies were then replica plated onto SC-Arg-Ura-Leu plates to identify chromosome loss events. Chromosome loss rate was determined by dividing the number of Arg+ Ura+ Leu–cells by the number of experimental cells used. The 95% confidence interval of the median chromosome loss rate for each strain was calculated using at least 16 isolates.

## TABLES

**Table S1.** Cryo-EM data collection, refinement, and validation.

|  | CTF3C-CNN1-WIP1<br>(DATASET 1) | CTF3C-CNN1-WIP1<br>(DATASET 2) |
| --- | --- | --- |
| **Data collection and processing** |  |  |
| Magnification | 105,000 | 105,000 |
| Voltage (kV) | 300 | 300 |
| Micrographs | 14,732 | 12,780 |
| Movies per stage position | 45 | 20 |
| Electron exposure (e-/Å<sup>2</sup>) | 52 | 52 |
| Exposure time (sec.) | 2.4 | 2.4 |
| Number of frames | 48 | 48 |
| Defocus range (μm) | -1.2 to -3.0 | -1.2 to -3.0 |
| Pixel size (Å) | .825 | .825 |
| Initial particle images (no.) | 5,673,375 | 2,963,710 |
| After 2D classification (no.) | 767,816 | 733,101 |
| Symmetry imposed | None | None |
| Final particle images (no.) | 96,787 |  |
| Map resolution (Å)<br>FSC threshold | 3.67<br>(0.143) |  |
| **Refinement** |  |  |
| Initial model used (PDB code) | 6WUC |  |
| Model resolution (Å)<br>FSC threshold | 3.9<br>(0.5) |  |
| Model resolution range (Å) | 40-3.7 |  |
| Map sharpening *B* factor (Å<sup>2</sup>) | NA |  |
| Model composition<br>Non-hydrogen atoms<br>Protein residues<br>Ligands | 5619<br>687<br>0 |  |
| *B* factors (Å<sup>2</sup>) | Min: 30.0<br>Max: 283.7<br>Mean: 177.31 |  |
| R.m.s. deviations<br>Bond lengths (Å)<br>Bond angles (°) | 0.005<br>0.899 |  |
| Validation<br>MolProbity score<br>Clashscore<br>Poor rotamers (%) | 2.42<br>23.49<br>0.16 |  |
| Ramachandran plot<br>Favored (%)<br>Allowed (%)<br>Disallowed (%) | 89.54<br>10.16<br>0.29 |  |

**Table S2.** Yeast strains used in this work.

| STRAIN | GENOTYPE | REFERENCE | FIGURE |
| --- | --- | --- | --- |
| HZY1203 | *MAT alpha CTF3-FLAG-ProtA:: HisMX*, W303 | This study | 3AC |
| HZY1202 | *MAT alpha ctf3-2A-FLAG-ProtA:: HisMX*, W303 | This study | 3AC |
| HZY600 | *MAT alpha LEU2 ura3-52 leu2Δ1 trp1Δ63 his3Δ200 lys2ΔBgl hom3-10 ade2Δ1 ade8* | Suhandynata *et al.*, 2019 | 3B,5B |
| HZY1265 | *ctf3-2A::HisMX*, derived from HZY600 | This study | 3B |
| HZY1622 | *ULP2-TAF::KanMX*, derived from HZY600 | This study | 3B |
| HZY1623 | *ulp2-KIM3A-TAF::KanMX*, derived from HZY600 | This study | 3B |
| HZY1624 | *ULP2-TAF::KanMX ctf3-2A::HisMX*, derived from HZY600 | This study | 3B |
| HZY1625 | *ulp2-KIM3A-TAF::KanMX ctf3-2A::HisMX*, derived from HZY600 | This study | 3B |
| HZY1255 | *MATa MCM21-ProtA::KanMX ulp2Δ::NatMX pRS316-ULP2*, W303 | This study | 3D |
| HZY1551 | *MATa MCM21-ProtA::KanMX ctf3-2A::HisMX ulp2Δ::NatMX pRS316-ULP2*, W303 | This study | 3D |
| HZY1256 | *MATa CTF3-FLAG-ProtA:: HisMX sml1Δ::TRP1 arg4Δ lys2ΔBgl* | This study | 4C |
| HZY1257 | *MATa ctf3-2A-FLAG-ProtA:: HisMX sml1Δ::TRP1 arg4Δ lys2ΔBgl* | This study | 4C |
| HZY632 | *MAT alpha ulp2-SIM<sup>3A</sup>:: kanMX*, derived from HZY600 | Suhandynata *et al.*, 2019 | 5A |
| HZY633 | *MAT alpha ulp2-KIM<sup>3A</sup>:: kanMX*, derived from HZY600 | Suhandynata *et al.*, 2019 | 5A |
| HZY607 | *MAT alpha ulp2- SIM<sup>3A</sup>KIM<sup>3A</sup>:: kanMX*, derived from HZY600 | Suhandynata *et al.*, 2019 | 5A |
| HZY601 | *MAT alpha arg4Δ::URA3 ura3-52 leu2Δ1 trp1Δ63 his3Δ200 lys2ΔBgl hom3-10 ade2Δ1 ade8* | Suhandynata *et al.*, 2019 | 5B |
| HZY1265 | *MAT alpha ctf3-2A::HisMX*, derived from HZY600 | This study | 5AB |
| HZY1269 | *MAT alpha ctf3-2A::HisMX ulp2-SIM<sup>3A</sup>:: kanMX*, derived from HZY600 | This study | 5AB |
| HZY1274 | *MAT alpha ctf3-2A::HisMX ulp2-KIM<sup>3A</sup>:: kanMX*, derived from HZY600 | This study | 5AB |
| HZY1267 | *MAT alpha ctf3-2A::HisMX ulp2-SIM<sup>3A</sup>KIM<sup>3A</sup>:: kanMX*, derived from HZY600 | This study | 5AB |
| HZY602 | *MAT alpha mad1Δ::HisMX*, derived from HZY600 | Suhandynata *et al.*, 2019 | 5A |
| HZY741 | *MATa sml1Δ::TRP1 rad53Δ::His3MX*, S288c | Suhandynata *et al.*, 2019 | 5A |
| HZY200 | SK1 diploid, *ura3/ura3 leu2/leu2 trp1/trp1 his3/his3 lys2/lys2 ade2/ADE2, arg4/ARG4, ho-del::LYS2/ho-del::LYS2* | This study | 5C |
| HZY222 | SK1 diploid, *ctf3-2A::HisMX/ctf3-2A::HisMX* | This study | 5C |
| HZY223 | SK1 diploid, *ulp2-SIM<sup>3A</sup>:: kanMX/ulp2-SIM<sup>3A</sup>:: kanMX* | This study | 5C |
| HZY224 | SK1 diploid, *ulp2-KIM<sup>3A</sup>:: kanMX/ulp2-KIM<sup>3A</sup>:: kanMX* | This study | 5C |
| HZY225 | SK1 diploid, *ulp2- SIM<sup>3A</sup>KIM<sup>3A</sup>:: kanMX/ulp2-SIM<sup>3A</sup>KIM<sup>3A</sup>:: kanMX* | This study | 5C |
| HZY231 | SK1 diploid, *ctf3-2A::HisMX/ctf3-2A::HisMX ulp2-SIM<sup>3A</sup>:: kanMX/ulp2-SIM<sup>3A</sup>:: kanMX* | This study | 5C |
| HZY3658 | *MATa ulp2Δ::NatMX pRS316-ULP2* | Suhandynata *et al.*, 2019 | S4A |
| SMH690 | *MATa MCM22-GFP::HisMX SPC110-mCherry::hphMX CTF3-3FLAG::KanMX* | Hinshaw *et al.*, 2019 | 4A |
| SMH740 | *MATa MCM22-GFP::HisMX SPC110-mCherry::hphMX ctf3Δ::KanMX* | This study | 4A |
| SMH738 | *MATa MCM22-GFP::HisMX SPC110-mCherry::hphMX CTF3-3FLAG::KanMX* | This study | 4A |
| SMH320 | *MATa SCC2-GFP::HisMX MTW1-tdTomato::natMX* | Hinshaw *et al.*, 2017 | S3A |
| SMH354 | *MATa SCC2-GFP::HisMX MTW1-tdTomato::natMX ctf19-9A::KanMX* | Hinshaw *et al.*, 2017 | S3A |
| SMH868 | *MATa SCC2-GFP::HisMX MTW1-tdTomato::natMX ctf3-2A::KanMX* | This study | S3A |
| SMH77 | *MATa SGO1-GFP::HisMX SPC110-mCherry::hphMX* | This study | S3B |
| SMH755 | *MATa SGO1-GFP::HisMX SPC110-mCherry::hphMX ctf19Δ::KanMX* | This study | S3B |
| SMH766 | *MATa SGO1-GFP::HisMX SPC110-mCherry::hphMX ctf3-2A::KanMX* | This study | S3B |

**Table S3.** Plasmids used in this work.

| PLASMID | GENOTYPE | REFERENCE | FIGURE |
| --- | --- | --- | --- |
| pSMH145 | pLIC-Tra His6-TEV-Ctf3; His6-TEV-Mcm16; His6-TEV-Mcm22 | Hinshaw et al., 2017 | 1B,S2A-B |
| pHZE1809/<br>pSMH1614 | pLIC-Tra His6-GST-TEV-Ulp2-896-1034 | Suhandynata, Quan, et al. 2019 | 1B,S1A-B,S2A |
| pHZE1810/<br>pSMH1615 | pLIC-Tra His6-GST- TEV-Ulp2-896-1034-3A | Suhandynata, Quan, et al. 2019 | 1B,S2A |
| pSMH1193 | pLICBac Cnn1 | Hinshaw and Harrison, 2019 | 1C |
| pSMH1198 | pLICBac His6-TEV-Wip1 | Hinshaw and Harrison, 2019 | 1C |
| pSMH1898 | pLIC-Tra His6-GST-TEV-Ulp2-SIM (717-734) | This study | S1A |
| pSMH1723 | pLIC-Tra His6-GST-TEV-Ulp2-907-937 | This study | S1B |
| pSMH1724 | pLIC-Tra His6-GST-TEV-Ulp2-917-937 | This study | S1B |
| pSMH1725 | pLIC-Tra His6-GST-TEV-Ulp2-927-937 | This study | S1B,2B-C |
| pSMH1711 | pLIC-Tra His6-TEV-Ctf3-2A; His6-TEV-Mcm16; His6-TEV-Mcm22 | This study | 2B,S2A |
| pSMH1727 | pLIC-Tra His6-GST-TEV-Ulp2-927-937-3A | This study | 2B-C |
| pSMH1760 | pLIC-Tra His6-GST-TEV-Ulp2-927-937-3DA | This study | 2C |
| pSMH1761 | pLIC-Tra His6-GST-TEV-Ulp2-927-937-1DA-1 | This study | 2C |
| pSMH1762 | pLIC-Tra His6-GST-TEV-Ulp2-927-937-1DA-2 | This study | 2C |
| pSMH1763 | pLIC-Tra His6-GST-TEV-Ulp2-927-937-1DA-3 | This study | 2C |
| pSMH1783 | pLIC-Tra His6-GST-TEV-Ulp2-927-937-1A-1 | This study | 2C |
| pSMH1784 | pLIC-Tra His6-GST-TEV-Ulp2-927-937-1A-2 | This study | 2C |
| pSMH1785 | pLIC-Tra His6-GST-TEV-Ulp2-927-937-1A-3 | This study | 2C |
| pSMH1711 | pLIC-Tra His6-TEV-Ctf3; His6-TEV-Mcm16; His6-TEV-Mcm22-2A-2 | This study | S2A |
| pSMH1728 | pLIC-Tra His6-GST-TEV-Ulp2-927-937-NA | This study | S2B |
| HZE2617 | pRS315-ULP2::kanMX | Suhandynata *et al.*, 2019 | 3D,S4A |
| HZE2618 | pRS315-ulp2-SIM<sup>3A</sup>::kanMX | Suhandynata *et al.*, 2019 | 3D,S4A |
| HZE2619 | pRS315-ulp2-SIM<sup>3A</sup>KIM<sup>3A</sup>::kanMX | Suhandynata *et al.*, 2019 | 3D,S4A |
| HZE2620 | pRS315-ulp2-KIM<sup>3A</sup>::kanMX | Suhandynata *et al.*, 2019 | 3D |
| HZE2902 | pRS315-ulp2-SIM<sup>3A</sup>D930A::kanMX | This study | S4A |
| HZE2903 | pRS315-ulp2-SIM<sup>3A</sup>V931A::kanMX | This study | S4A |
| HZE2904 | pRS315-ulp2-SIM<sup>3A</sup>L933A::kanMX | This study | S4A |
| HZE2905 | pRS315-ulp2-SIM<sup>3A</sup>I934A::kanMX | This study | S4A |

